# Evaluation of Correlation between CD44, Radiotherapy Response and Survival Rate in Patients with Advanced Stage of Head and Neck Squamous Cell Carcinoma (HNSCC)

**DOI:** 10.1101/2021.09.26.461653

**Authors:** Parul Dubey, Rajeev Gupta, Anupam Mishra, Vijay Kumar, Smrati Bhadauria, Madan Lal Brahma Bhatt

## Abstract

**Purpose:** Cancer stem cells (CSCs) constitute a distinctive subpopulation of cancer cells that are competent in tumor initiation, invasion, recurrence, and resistance to chemo-radiotherapy. CD44, a hyaluronic acid (HA) receptor has been considered as a potential CSC marker in head and neck cancer. The purpose of this study is to evaluate the correlation between CD44 and clinicopathological parameters, treatment response, survival, and recurrence.

**Methods:** CD44 expression was examined by immunohistochemistry (IHC) in 90 samples of head and neck squamous cell carcinoma (HNSCC) confirmed patients. Expression of CD44 and its association with clinicopathological parameters, treatment response, and survival was determined.

**Results:** In all HNSCC patient samples, CD44 was expressed consistently at different intensities. Tumor size (p<0.001), stage (p<0.001) and treatment response (p<0.001) showed statistically significant association with CD44 expression. Alcohol and CD44 were observed as independent predictors of response to radiotherapy by using multivariate ordinal logistic regression analysis. Analysis of 2 years overall survival (OS) showed that CD44 expression (p=0.02), tumor size (p=0.001), lymph node status (p<0.001), stage (p<0.001) and grade (p=0.007) were significantly associated with OS. By using Cox regression analysis, lymph node status (p=0.001), grade (p<0.001), recurrence (p<0.001) and CD44 expression (p=0.003) were found to be potential independent predictors of OS.

**Conclusion:** Our findings suggest that CD44 contributes to resistance to radiotherapy and poor OS. The results also suggest that except for CD44 there could be other factors such as lymph node metastasis, grade, and alcohol which should be investigated as potential targets for therapy.

## Introduction

Head and neck squamous cell carcinoma (HNSCC) is the sixth most common cancer worldwide affecting over 400,000 patients annually leading to over 200,000 deaths^1^. Although advent of newer surgical and endoscopic techniques over last 30-40 years have improved the quality of life^2^, however, the 5 year survival rate has still remained same as earlier (40-50%)^3^. Despite the treatments such as surgery and/or combination of chemotherapy and radiotherapy, the prognosis of HNSCC remains poor due to late stage diagnosis, high rates of primary site recurrence and common metastasis to loco-regional lymph nodes^4-8^. In case of HNSCC, patients with HPV positive disease exhibited poor prognosis. Especially in case of oropharynx cancer, HPV infection was considered as the major causative agent. In United States, it accounts for approximately 60-70% of oropharynx cancer. Within Western Europe, the prevalence varies ranging between 6.1 and 75% ^9^. However, in third world countries, HPV(+) HNSCC is relatively rare with a <10% prevalence^10^. Novel approaches towards increasing the overall and disease free survival as well as improved prognosis/patient outcome are much needed.

Cancer stem cells (CSCs) were first found in leukemia, and since then their presence has been confirmed in various other malignancies^11^. It has been postulated that CSC play a significant role in the pathobiology of HNSCC^12^. CSCs are cancer cells having capacity for self renewal, differentiation and regeneration. Recent studies have demonstrated that CSCs are found within the tumor and are resistant to current treatment modalities leading to relapse. Therefore, CSCs could be the potential therapeutic targets for the better treatment of HNSCC patients^13^.

HNSCCs have heterogeneous cellular composition. Various putative CSC markers reported to be present in HNSCC cell lines are ALDH, CD133, CD44 and others. CD44 was one of the very first CSC markers to be identified in HNSCC^14^. CD44 is a type I transmembrane glycoprotein, receptor for hyaluronic acid and functions as a major adhesion molecule^15^. It is involved in aggregation, proliferation, migration and angiogenesis. Adhesion to components of the extracellular matrix is influenced by the interaction between hyaluronic acid and CD44^16^. It has been reported that high levels of Bmi-1 gene was expressed in CD44+ cells^14^ and its role has been found in self-renewal and tumorigenesis^17, 18^.

Various cell line based studies have substantiated the presence of CSCs as one of the critical determinants of response/relapse of cancer cells to radiotherapy. However, definitive clinical investigations for assessing predictive significance of CSCs with reference to radiotherapy are lacking. In view of this, current study was planned to evaluate the expression of CSC marker "CD44" in relation to the clinicopathological features of HNSCC patients vis a vis their response to radiotherapy. Predictive significance of CD44 was assessed in determining response/relapse of HNSCC patients to radiotherapy as well as clinical outcome and overall survival (OS).

## Materials and Methods

### Study Population

From March 2016 to March 2017, total 90 histopathologically confirmed patients for head and neck squamous cell carcinoma (HNSCC) were recruited in this study for the evaluation of cancer stem cell marker “CD44” by immunohistochemistry (IHC). All inoperable and non-metastatic patients strongly recommended only for chemo-radiotherapy treatment were included in the study. Patients with HPV infection, prior history of chemo-radiotherapy or surgery and distant metastasis were excluded from the study. Patients having habits of consuming tobacco and alcohol were included. Patients consuming alcohol are categorized into three categories: 1) former drinker, 2) occasional drinker (consumed alcohol once in a week) and 3) heavy drinkers (consumed alcohol daily). These patients underwent biopsy. Healthy matched individuals (n=90) served as control group, attending dental procedures for dental implant, tooth extraction or benign cyst. The HNSCC tissues were extracted from various sites of oral cavity and neck region. All the tissues collected were stored in formalin at room temperature. Only pre-treated tissues were used to measure the CD44 expression by IHC. The Institutional Ethics Committee has approved the study protocol, which follows the Declaration of Helsinki. The written informed consent has been obtained from all the patients included in this study.

### Treatment and Response Assessment

HNSCC patients were treated with standard treatment of Radiotherapy. Radiotherapy was done by using linear accelerator. For 7 weeks, a dose of 70Gy/35 fractions, 5 Fc/week of radiation was delivered (46Gy/23# to primary tumor and whole neck followed by 24Gy/12# to tumor and neck sparing the cord). For 7 weeks, 30mg/m^2^ dose of cisplatin with a ceiling of absolute dose of 50mg i.v. weekly was given to fit and adult patients. All patients had received concurrent chemo-radiotherapy and cisplatin was given to all the patients. On the basis of 30mg/m^2^ cispaltin dose, all patients had received 210mg/m^2^ total dose of cisplatin for 7 weeks. Response was seen after one month of completion of treatment. The characteristics of HNSCC patients which includes age, clinical stage, TNM classification (defined by the American Joint Committee on Cancer, AJCC 2010, VII Edition)^19^ and oral habits (Tobacco and alcohol intake) were assessed by radiation oncologist. Assessment of tumor response was done by standard clinical examination and radiological investigation. The treatment response was stratified on the basis of guidelines defined by Response Evaluation Criteria in Solid Tumors (RECIST) which is a tumor-centric set of published rules that define whether the tumor regresses (responsive), stay the same (stabilize) or worsen (progress) during the treatment. The response criteria included complete response (CR), i.e., disappearance of all target lesions and reduction of tumor in short axis to <10mm; partial response (PR), i.e., reduction of at least 30% in the diameter of target lesion and no response (NR) [stable disease (SD) + progressive disease (PD)]. Progressive disease, i.e., increase by at least 20% of diameter of target lesion or emergence of one or more new lesions; stable disease, i.e., inadequate shrinkage of tumor which meet the requirements for PR or inadequate increase in the size of tumor which meet the requirements for PD^20^.

After treatment, patients were followed up and evaluated for survival. Treatment response and overall survival (OS) starting from the date of diagnosis to the date of death were considered as the primary endpoints. After the treatment and evaluation, response to radiotherapy was correlated with CD44 expression to find out the role of CSCs as a predictive biomarker of radiotherapy response.

### Immunohistochemistry

Immunohistochemical detection of CSC marker, CD44 also known as HCAM was performed on a formalin-fixed, paraffin-embedded tissue sections (5μm). HNSCC tissue slides were deparaffinized, rehydrated and washed. Endogenous peroxidases were blocked by using 0.1% hydrogen peroxide for 15min at room temperature. Prior to this heat-induced epitope retrieval (HIER) with citrate buffer (pH 6.0) was done for 30min. The deparaffinized and rehydrated tissue sections were incubated overnight with the primary antibody, viz., anti-CD44 mouse monoclonal antibody [HCAM (DF-1485): sc-7297 Santa Cruz Biotechnology, California, USA], at 1:50 dilution. Thereafter, samples were incubated for 1hr with HRP-conjugated goat anti mouse IgG secondary antibody at 1:200 dilutions. Antigen was visualized with DAB-Peroxidase Substrate (Thermo Fisher Scientific, Waltham, MA, USA). Finally, tissue specimens were stained with hematoxylin (Thermo Fisher Scientific, USA) to discriminate nucleus from cytoplasm. Thereafter, sections were mounted in DPX (Sigma) and analyzed at 40X magnification using Lieca DCF450C bright field/fluorescence microscope. At least two different sections per sample were analyzed. Whole tissue section was evaluated and 10 different fields per section were observed for positive staining. It was grouped as following: No staining (0%), Low staining (≤50%) and high staining (>50%)^21^.

### Statistical Analysis

Continuous data were summarized as Mean ± SE while discrete (categorical) in number (n) and percentage (%). Continuous groups were compared by independent Student"s t test. Categorical groups were compared by chi-square (χ^2^) test. Univariate and multivariate ordinal logistic regression analysis was done to assess independent predictor(s) of responsiveness to radiotherapy. Survival analysis was done using Kaplan-Meier method and difference between groups was done by Log-rank test. A two-tailed (*α*=2) p<0.05 was considered statistically significant. Analysis was performed on GraphPad Prism 5.01 and IBM SPSS Statistics 20 software.

## Results

### Basic Characteristics of HNSCC Patients

The basic characteristics (demographic, clinico-pathological and treatment response) of HNSCC patients are summarized in Table 1. The mean age (±SE) of patients was 46.64 ± 1.19 years and median was 45 years. Most of the patients were of age ≤ 45 years (54.4%) and majority was males (86.7%). Among all the 90 patients, patients who consumed tobacco in any form were 76.7%, and patients who consumed alcohol were 41.1%.

**Table 1:**
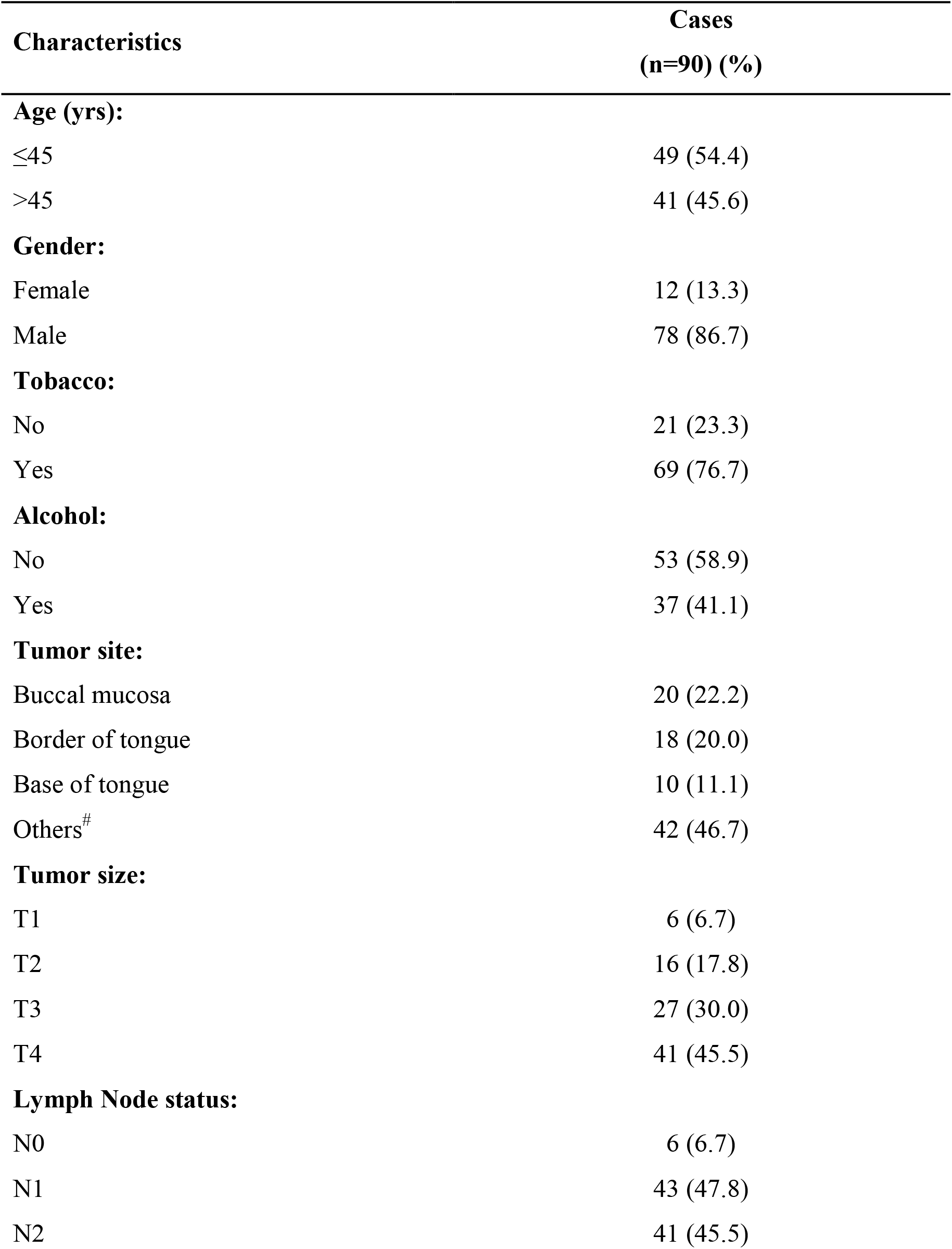
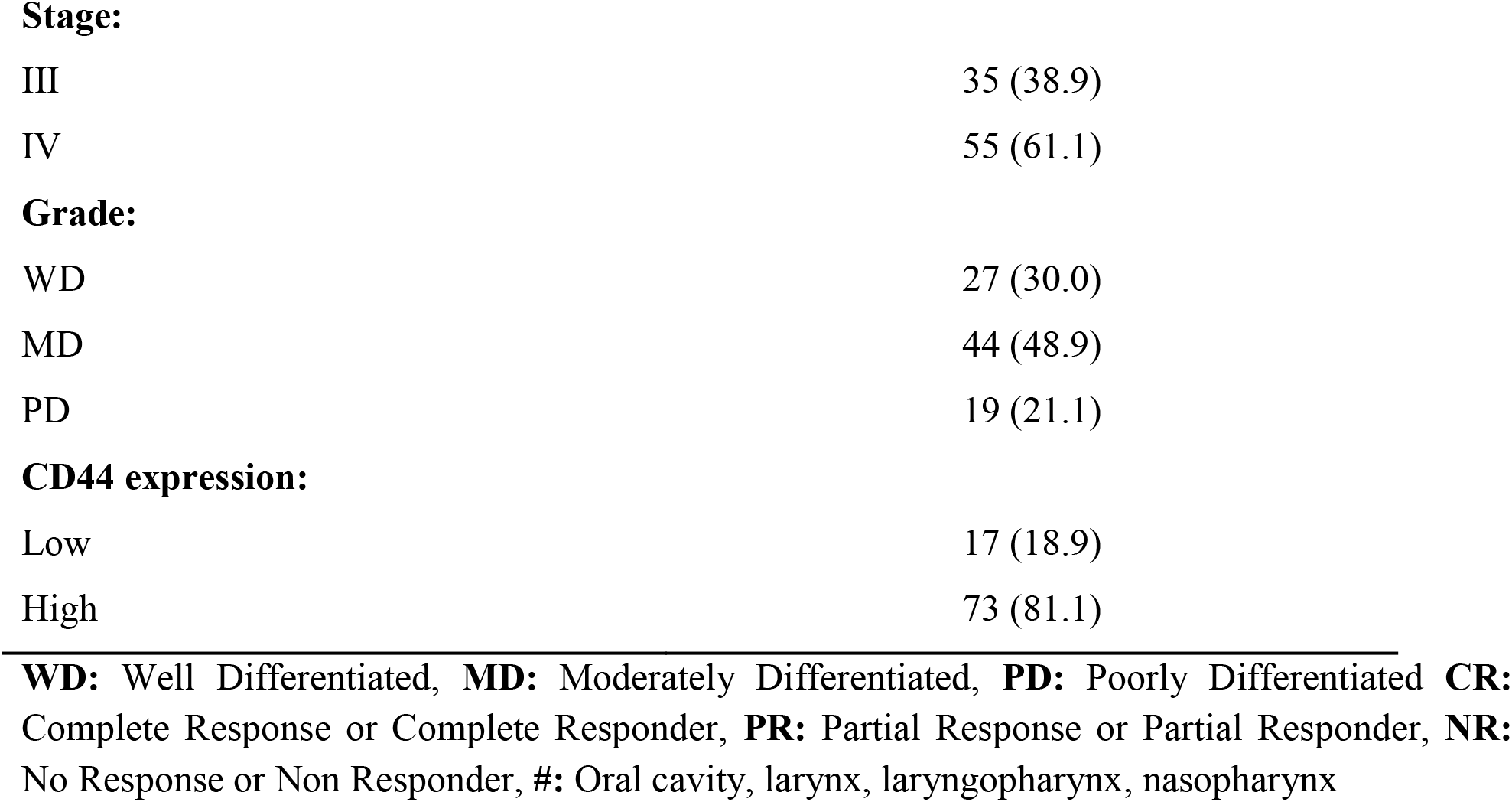
Demographic, clinico-pathological characteristics and molecular marker of HNSCC patients.

Buccal mucosa 20 (22.2%) was the primary tumor site in majority of the patients (Suppl 1). Furthermore, on considering clinico-pathological parameters it was observed that patients were mostly having T4 tumor size (45.5%), N1 lymph node (47.8%), IV stage (61.1%) and moderately differentiated (MD) grade (48.9%).

On taking treatment response into account, 19 (21.1%) patients had complete response (CR), 20 (22.2%) patients had partial response (PR) and 51 (56.7%) had no response (NR). Therefore, the overall response (CR+PR) was 43.3%.

### Assessment of Molecular Marker Expression

The expression of molecular marker CD44 was observed by immunohistochemistry shown in Fig. 1 (A, B). CD44 marker was positively expressed in all the patients. However, 17 (18.9%) patients showed low expression and 73 (81.1%) had high expression. Majority of tumor specimens showing CD44 expression were collected from oral cavity (Suppl 2). No staining (0%) for CD44 was observed in control tissue specimens. Immunohistochemical detection of CD44 specimens in HNSCC patients revealed that at cellular periphery CD44 was predominantly localized. Localization of CD44 was also observed prominently at the invasive front in all HNSCC specimens.

**Fig 1:**
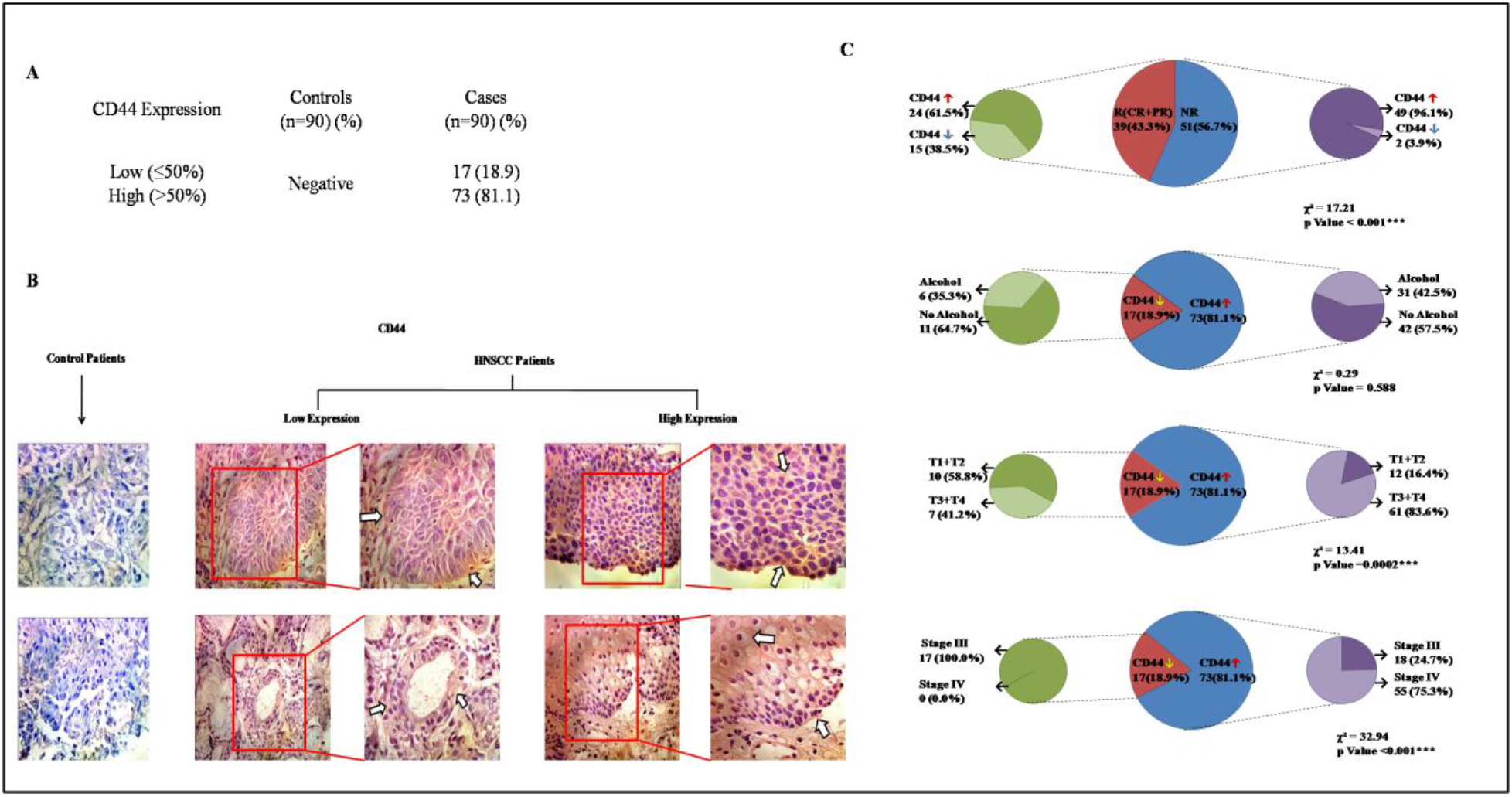
Immunohistochemistry detection of CD44 in HNSCC specimens and pie chart depicting the association between various variables using Chi-square test. **A:** Frequency distribution of CD44 expression in control and HNSCC patients. **B:** Micrograph showing negative staining from control samples. Positive signals were obtained in all the HNSCC tissue specimens showing low and high expression of positively stained foci. The CD44 was predominantly localized at the cellular periphery. A prominent localization of CD44 was observed mainly in the HNSCC cells present at the invasive front (arrow). **C:** Showing significant association between treatment response and CD44 expression (p<0.001); Showing significant association between CD44 expression and tumor size (p=0.0002) and stage (p<0.001); Showing no significant association between CD44 expression and alcohol (p=0.588).

### Association of Treatment Response with Demographic, Clinico-pathological Parameters and Molecular marker Expression

An association was established between treatment response and various variables (demographic, clinico-pathological parameters) which are summarized in Table 2. The significant association was observed between treatment response and alcohol (χ^2^ = 6.48, p = 0.039), tumor size (χ^2^ = 13.26, p = 0.039) and stage (χ^2^ = 8.85, p = 0.012). Significant association between treatment response and CD44 expression (χ^2^ = 17.21, p<0.001) was observed which is presented in Fig. 1 (C). These significant associations suggest that these variables could serve as the predictors of responsiveness.

**Table 2:**
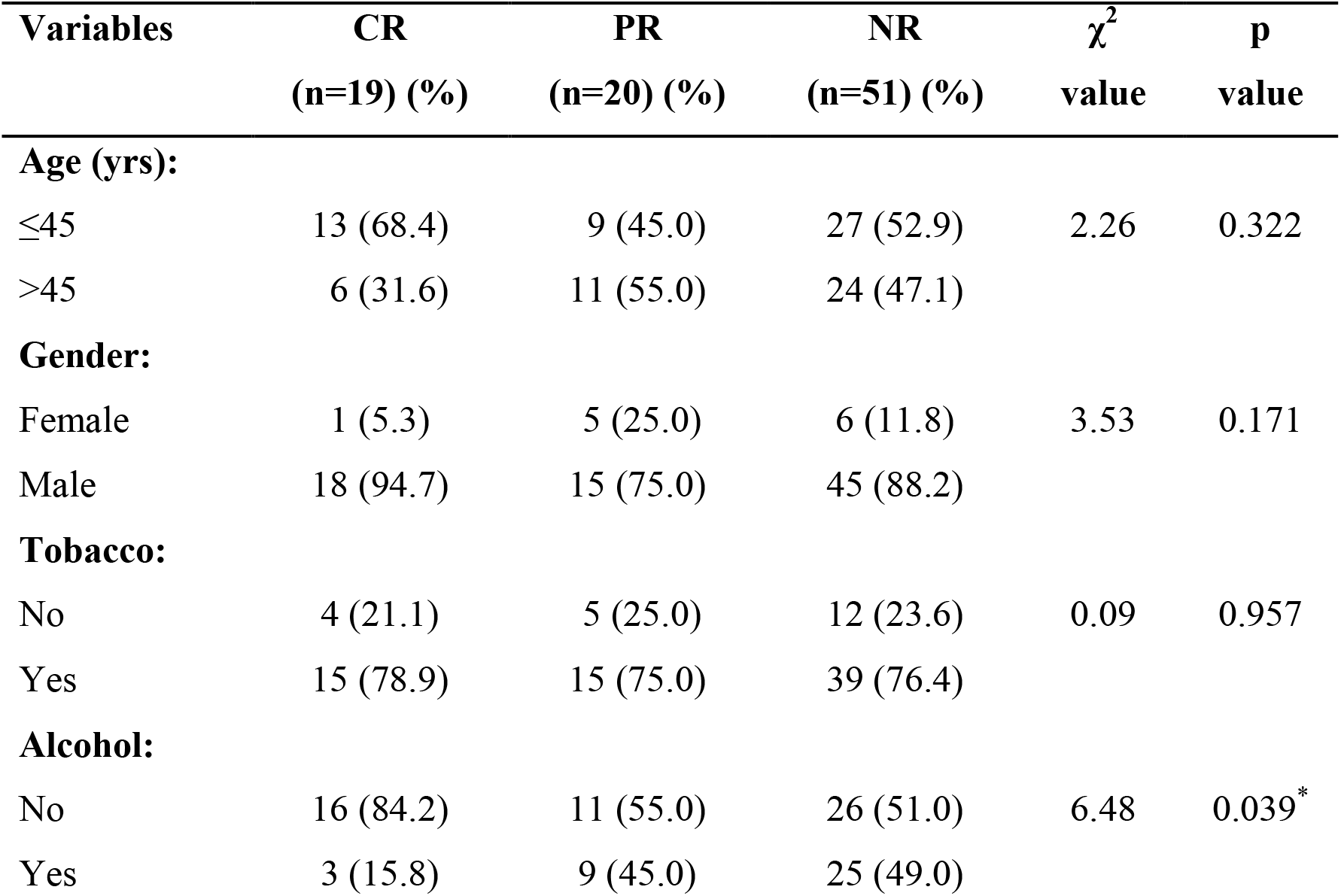
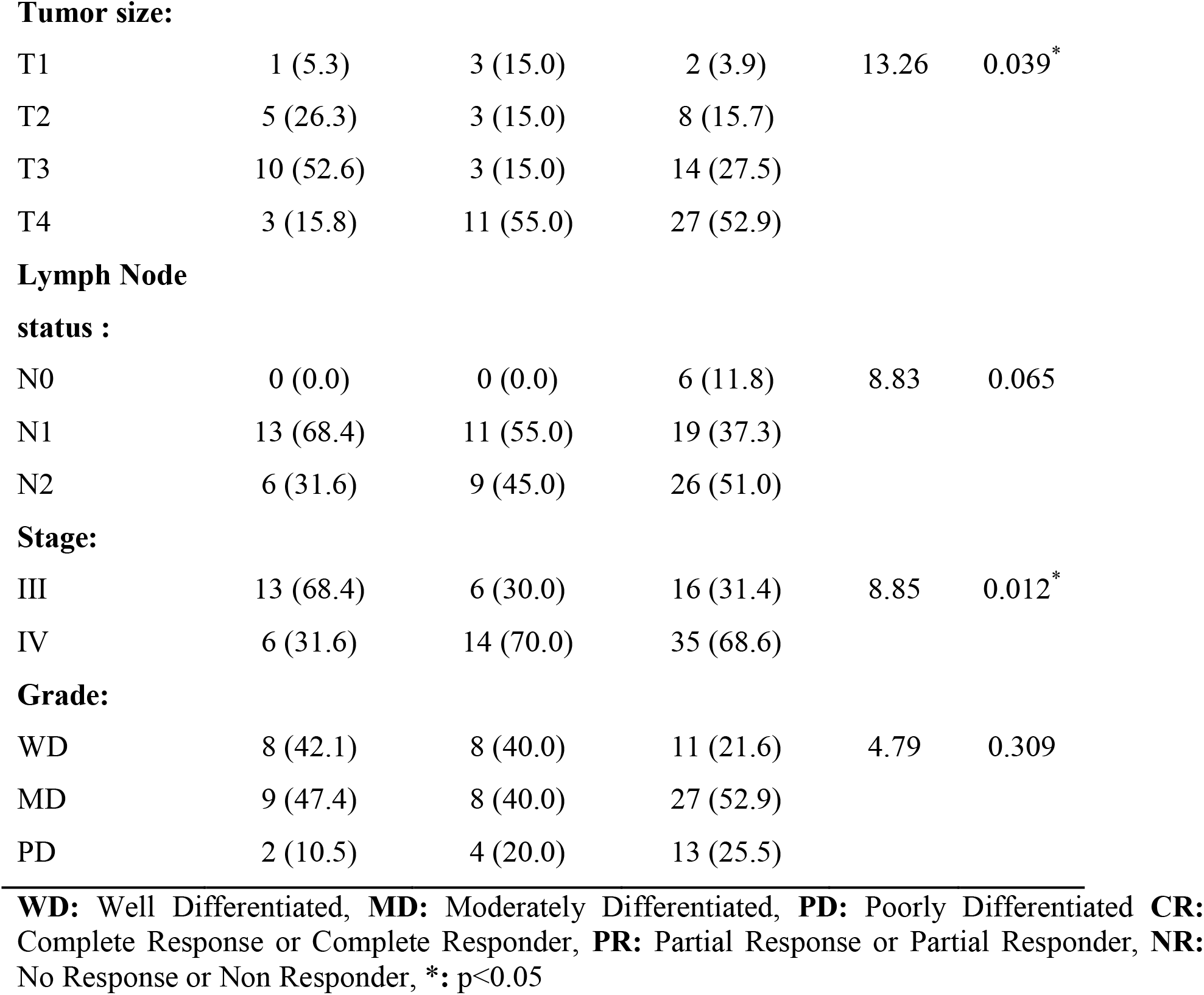
Association of treatment response with demographic, clinical and biomarker expression of patients (n=90)

| Variables | CR<br>(n=19) (%) | PR<br>(n=20) (%) | NR<br>(n=51) (%) | $\chi^2$<br>value | p<br>value |
| --- | --- | --- | --- | --- | --- |
| **Age (yrs):** |  |  |  |  |  |
| ≤45 | 13 (68.4) | 9 (45.0) | 27 (52.9) | 2.26 | 0.322 |
| >45 | 6 (31.6) | 11 (55.0) | 24 (47.1) |  |  |
| **Gender:** |  |  |  |  |  |
| Female | 1 (5.3) | 5 (25.0) | 6 (11.8) | 3.53 | 0.171 |
| Male | 18 (94.7) | 15 (75.0) | 45 (88.2) |  |  |
| **Tobacco:** |  |  |  |  |  |
| No | 4 (21.1) | 5 (25.0) | 12 (23.6) | 0.09 | 0.957 |
| Yes | 15 (78.9) | 15 (75.0) | 39 (76.4) |  |  |
| **Alcohol:** |  |  |  |  |  |
| No | 16 (84.2) | 11 (55.0) | 26 (51.0) | 6.48 | 0.039* |
| Yes | 3 (15.8) | 9 (45.0) | 25 (49.0) |  |  |
| **Tumor size:** |  |  |  |  |  |
| T1 | 1 (5.3) | 3 (15.0) | 2 (3.9) | 13.26 | 0.039* |
| T2 | 5 (26.3) | 3 (15.0) | 8 (15.7) |  |  |
| T3 | 10 (52.6) | 3 (15.0) | 14 (27.5) |  |  |
| T4 | 3 (15.8) | 11 (55.0) | 27 (52.9) |  |  |
| **Lymph Node****status :** |  |  |  |  |  |
| N0 | 0 (0.0) | 0 (0.0) | 6 (11.8) | 8.83 | 0.065 |
| N1 | 13 (68.4) | 11 (55.0) | 19 (37.3) |  |  |
| N2 | 6 (31.6) | 9 (45.0) | 26 (51.0) |  |  |
| **Stage:** |  |  |  |  |  |
| III | 13 (68.4) | 6 (30.0) | 16 (31.4) | 8.85 | 0.012* |
| IV | 6 (31.6) | 14 (70.0) | 35 (68.6) |  |  |
| **Grade:** |  |  |  |  |  |
| WD | 8 (42.1) | 8 (40.0) | 11 (21.6) | 4.79 | 0.309 |
| MD | 9 (47.4) | 8 (40.0) | 27 (52.9) |  |  |
| PD | 2 (10.5) | 4 (20.0) | 13 (25.5) |  |  |

**WD:** Well Differentiated, **MD:** Moderately Differentiated, **PD:** Poorly Differentiated **CR:** Complete Response or Complete Responder, **PR:** Partial Response or Partial Responder, **NR:** No Response or Non Responder, \*: p<0.05

Since among all the demographic and clinico-pathological variables only alcohol, tumor size and stage showed the significant association, we next tried to establish an association of CD44 expression with all the above mentioned variables and it is presented in Fig. 1 (C).

Tumor size (χ^2^ = 13.41, p<0.0002) and stage (χ^2^ = 32.94, p<0.001) showed highly significant association with CD44 expression. On the other hand, alcohol (χ^2^ = 0.29, p = 0.588) was not found significantly associated with CD44 expression.

### Predictors of Radiotherapy Response

The predictor(s) of response (CR, PR and NR) to radiotherapy in HNSCC patients was determined by examining the patient"s demographic, clinico-pathological characteristics and CD44 expression using univariate and multivariate ordinal logistic regression analysis. It was considered that responsiveness is the dependent variable and demographic, clinico-pathological and CD44 expression are the independent variable and it is summarized in Table 3.

**Table 3:**
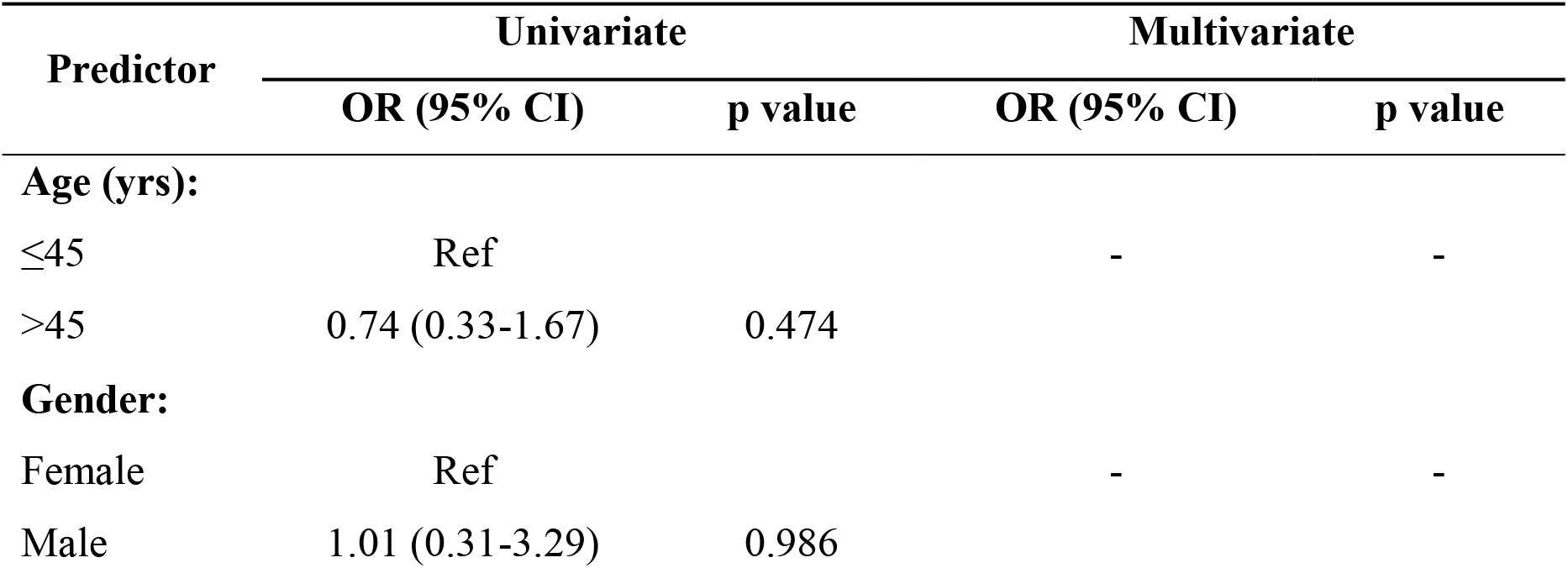
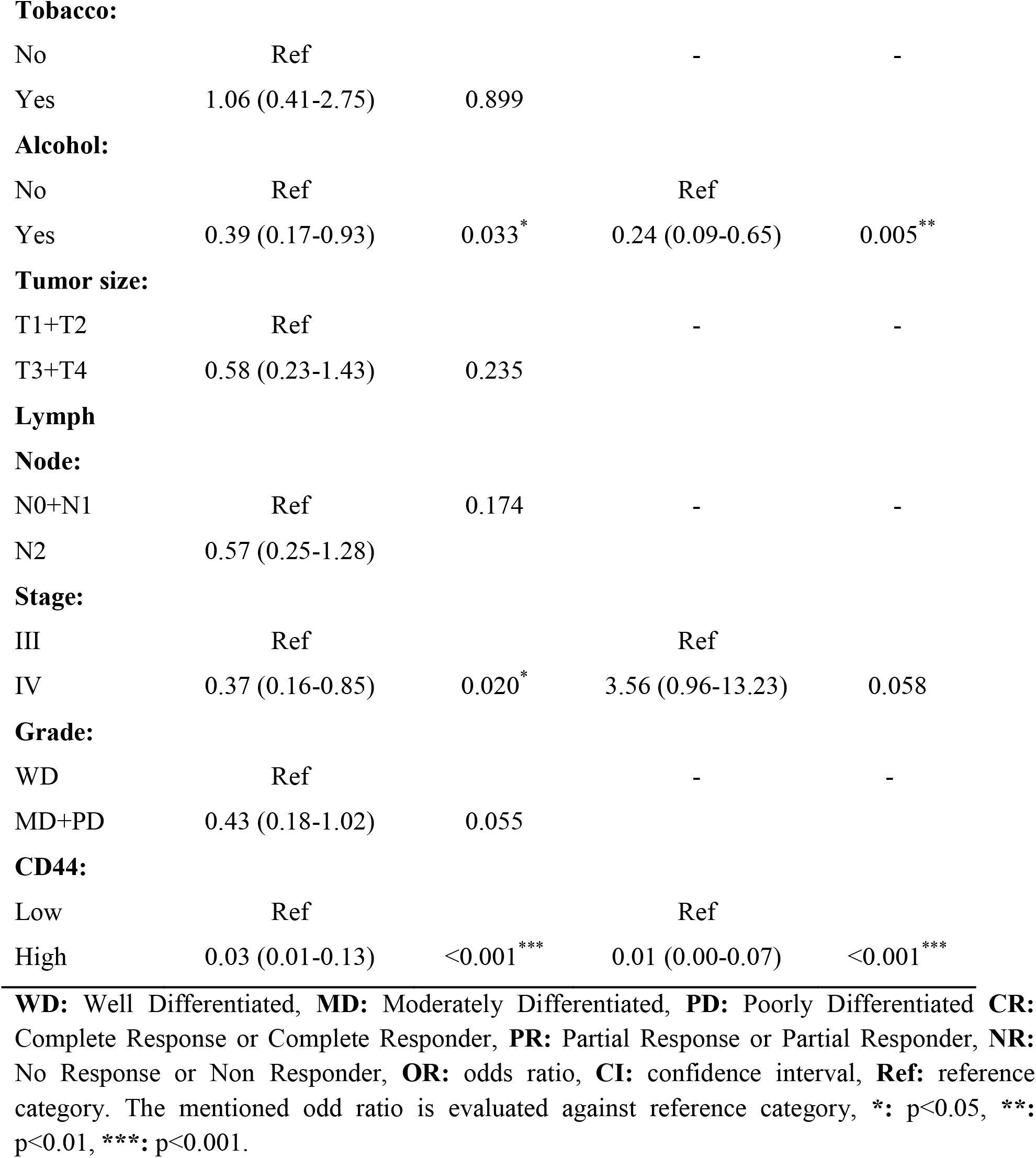
Identification of predictor of responsiveness to radiotherapy in HNSCC patients using ordinal logistic regression analysis (n=90)

| **Predictor** | **Univariate** |  | **Multivariate** |  |
| --- | --- | --- | --- | --- |
|  | **OR (95% CI)** | **p value** | **OR (95% CI)** | **p value** |
| **Age (yrs):** |  |  |  |  |
| ≤45 | Ref |  | - | - |
| >45 | 0.74 (0.33-1.67) | 0.474 |  |  |
| **Gender:** |  |  |  |  |
| Female | Ref |  | - | - |
| Male | 1.01 (0.31-3.29) | 0.986 |  |  |
| **Tobacco:** |  |  |  |  |
| No | Ref |  | - | - |
| Yes | 1.06 (0.41-2.75) | 0.899 |  |  |
| **Alcohol:** |  |  |  |  |
| No | Ref |  | Ref |  |
| Yes | 0.39 (0.17-0.93) | 0.033* | 0.24 (0.09-0.65) | 0.005** |
| **Tumor size:** |  |  |  |  |
| T1+T2 | Ref |  | - | - |
| T3+T4 | 0.58 (0.23-1.43) | 0.235 |  |  |
| **Lymph****Node:** |  |  |  |  |
| N0+N1 | Ref | 0.174 | - | - |
| N2 | 0.57 (0.25-1.28) |  |  |  |
| **Stage:** |  |  |  |  |
| III | Ref |  | Ref |  |
| IV | 0.37 (0.16-0.85) | 0.020* | 3.56 (0.96-13.23) | 0.058 |
| **Grade:** |  |  |  |  |
| WD | Ref |  | - | - |
| MD+PD | 0.43 (0.18-1.02) | 0.055 |  |  |
| **CD44:** |  |  |  |  |
| Low | Ref |  | Ref |  |
| High | 0.03 (0.01-0.13) | <0.001*** | 0.01 (0.00-0.07) | <0.001*** |

**WD:** Well Differentiated, **MD:** Moderately Differentiated, **PD:** Poorly Differentiated **CR:** Complete Response or Complete Responder, **PR:** Partial Response or Partial Responder, **NR:** No Response or Non Responder, **OR:** odds ratio, **CI:** confidence interval, **Ref:** reference category. The mentioned odd ratio is evaluated against reference category, \*: p<0.05, \*\*: p<0.01, \*\*\*: p<0.001.

In univariate ordinal logistic regression analysis, alcohol, stage and CD44 expression were found to be significantly associated with responsiveness.

Furthermore, when these significant predictors were subjected to multivariate analysis, alcohol and CD44 expression showed significant association with responsiveness. It indicates that alcohol and CD44 expression are independent predictors of responsiveness to radiotherapy in HNSCC patients.

### Overall Survival of HNSCC Patients and its Association with Clinico-pathological Characteristics, Biomarker Expression and Treatment Response

The HNSCC patients were followed up for 2 years after treatment. During this period 20 patients (22.2%) died due to disease, 53 (58.9%) patients remained alive while 17 (18.9%) were lost to follow up (LOF). All together this account for total 70 alive (Alive + LOF) patients, i.e., 77.8% was the incidence of overall survival. Further during the follow up, it was observed that 37 (41.1%) patients out of 90 patients showed recurrence.

Analysis of 2 year overall survival of HNSCC patients with clinico-pathological parameters, CD44 expression and treatment response was done by using Kaplan-Meier method. The difference between the groups was done by using Log rank (Mantel-Cox) test and it is summarized graphically in Fig. 2. With overall survival, all clinico-pathological characteristics and biomarker expression was found to be significantly (p<0.05 or p<0.01 or p<0.001) associated in our analysis using Log rank (Mantel-Cox) test. The Log rank test showed that mean survival decreases as the tumor size (χ^2^=16.34, p=0.001), lymph node (χ^2^=20.57, p<0.001), stage (χ^2^=14.03, p=0.0002) and grade (χ^2^=9.84, p=0.007) increases. In addition to this, it was observed that mean survival also decreases with high expression of CD44 (χ^2^=5.93, p=0.02) as compared to low expression of CD44. Moreover, our analysis showed that treatment response, i.e., CR, PR and NR was not associated significantly with overall survival (χ^2^=3.83, p=0.147).

**Figure 2:**
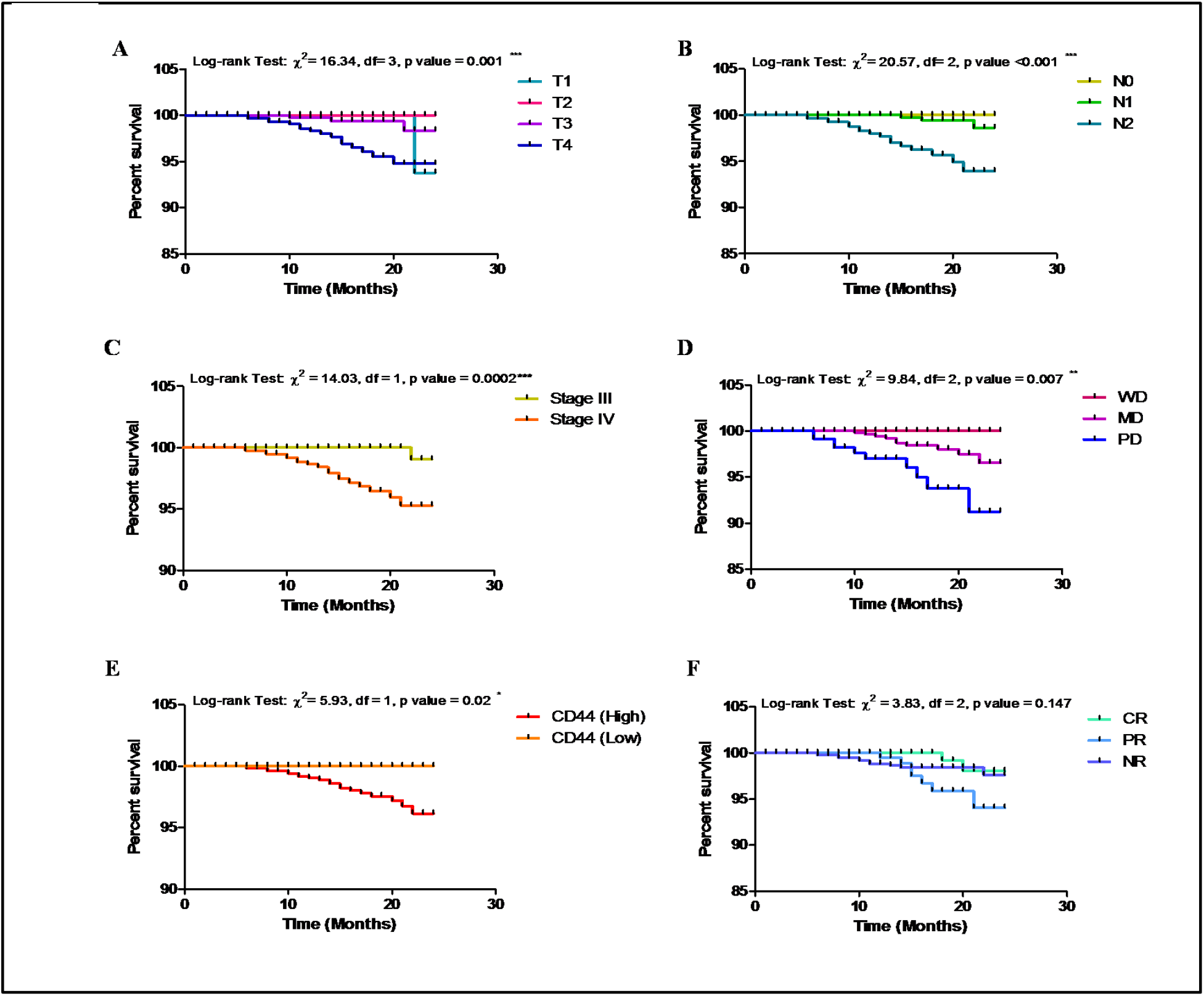
Two year overall survival of HNSCC patients using Kaplan-Meier method. **A:** Overall survival showing significant association with tumor size. **B:** Overall survival showing significant association with lymph node status. **C:** Overall survival showing significant association with stage. **D:** Overall survival showing significant association with grade. **E:** Overall survival showing significant association with CD44 expression. **F:** Overall survival showing no significant association with treatment response.

### Independent predictors of survival

Univariate and multivariate Cox regression analysis was done to determine the independent predictors of time dependent survival (death and live + LOF). Cox regression analysis was done between survival and different predictor variables (demographic, clinico-pathological, molecular marker expression, treatment response and recurrence) which are summarized in Table 4.

**Table 4:**
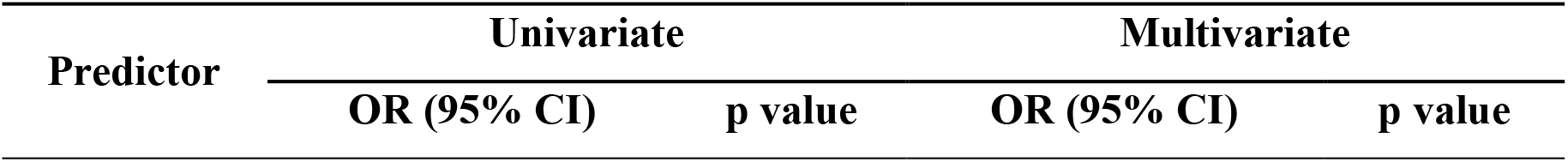
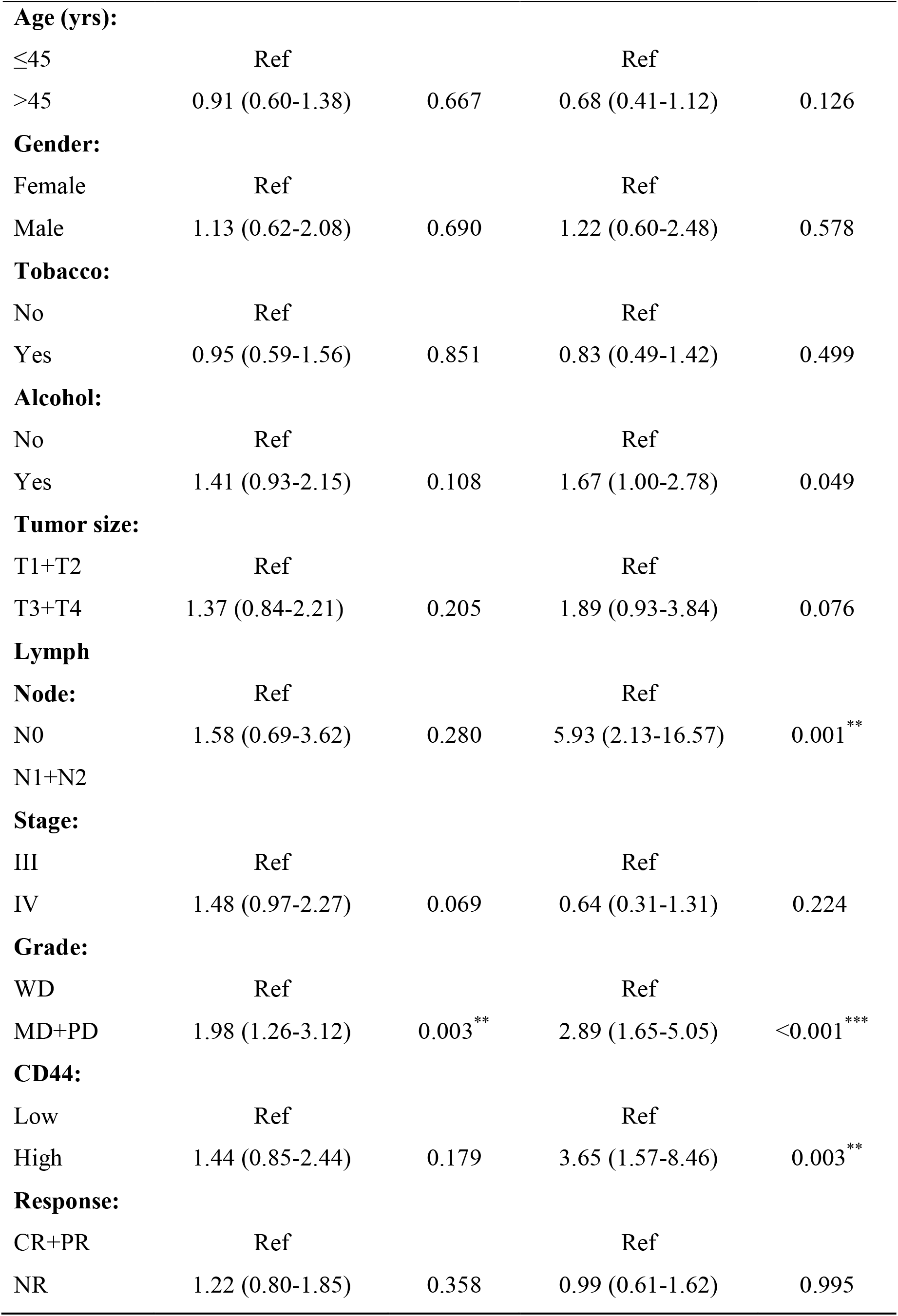
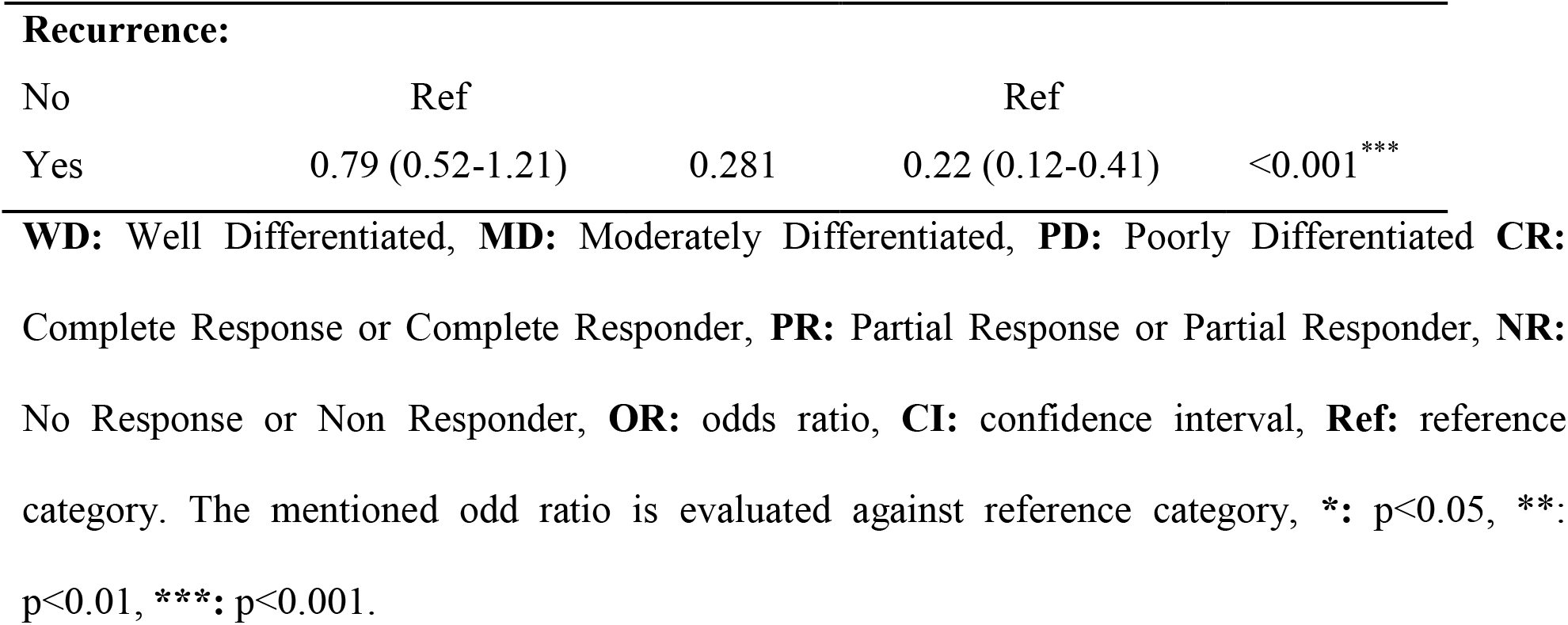
Time dependent independent predictors of survival using univariate and multivariate Cox regression analysis (n=90)

| Predictor | Univariate |  | Multivariate |  |
| --- | --- | --- | --- | --- |
|  | OR (95% CI) | p value | OR (95% CI) | p value |
| **Age (yrs):** |  |  |  |  |
| ≤45 | Ref |  | Ref |  |
| >45 | 0.91 (0.60-1.38) | 0.667 | 0.68 (0.41-1.12) | 0.126 |
| **Gender:** |  |  |  |  |
| Female | Ref |  | Ref |  |
| Male | 1.13 (0.62-2.08) | 0.690 | 1.22 (0.60-2.48) | 0.578 |
| **Tobacco:** |  |  |  |  |
| No | Ref |  | Ref |  |
| Yes | 0.95 (0.59-1.56) | 0.851 | 0.83 (0.49-1.42) | 0.499 |
| **Alcohol:** |  |  |  |  |
| No | Ref |  | Ref |  |
| Yes | 1.41 (0.93-2.15) | 0.108 | 1.67 (1.00-2.78) | 0.049 |
| **Tumor size:** |  |  |  |  |
| T1+T2 | Ref |  | Ref |  |
| T3+T4 | 1.37 (0.84-2.21) | 0.205 | 1.89 (0.93-3.84) | 0.076 |
| **Lymph Node:** |  |  |  |  |
|  | Ref |  | Ref |  |
| N0 | 1.58 (0.69-3.62) | 0.280 | 5.93 (2.13-16.57) | 0.001** |
| N1+N2 |  |  |  |  |
| **Stage:** |  |  |  |  |
| III | Ref |  | Ref |  |
| IV | 1.48 (0.97-2.27) | 0.069 | 0.64 (0.31-1.31) | 0.224 |
| **Grade:** |  |  |  |  |
| WD | Ref |  | Ref |  |
| MD+PD | 1.98 (1.26-3.12) | 0.003** | 2.89 (1.65-5.05) | <0.001*** |
| **CD44:** |  |  |  |  |
| Low | Ref |  | Ref |  |
| High | 1.44 (0.85-2.44) | 0.179 | 3.65 (1.57-8.46) | 0.003** |
| **Response:** |  |  |  |  |
| CR+PR | Ref |  | Ref |  |
| NR | 1.22 (0.80-1.85) | 0.358 | 0.99 (0.61-1.62) | 0.995 |
| **Recurrence:** |  |  |  |  |
| No | Ref |  | Ref |  |
| Yes | 0.79 (0.52-1.21) | 0.281 | 0.22 (0.12-0.41) | <0.001*** |

**WD:** Well Differentiated, **MD:** Moderately Differentiated, **PD:** Poorly Differentiated **CR:** Complete Response or Complete Responder, **PR:** Partial Response or Partial Responder, **NR:** No Response or Non Responder, **OR:** odds ratio, **CI:** confidence interval, **Ref:** reference category. The mentioned odd ratio is evaluated against reference category, \*: p<0.05, \*\*: p<0.01, \*\*\*: p<0.001.

In univariate analysis, only grade showed the significant association with the survival. However, in multivariate analysis, lymph node, grade, CD44 expression and recurrence showed significant association with the survival. Hence, it could be inferred that after adjusting the variables lymph node, grade, CD44 expression and recurrence could serve as the independent predictors of survival.

## Discussion

Cancer stem cells (CSCs), a small population of cancer cells, were revealed to be responsible for tumor initiation, relapse/recurrence and chemo or radiotherapy resistance^22,23^. In HNSCC, several cell surface markers have been reported as CSC marker such as CD44, CD133, ALDH1 and ABCG2^24-26^. These markers exhibiting high expression usually indicate poor prognosis. Among all these markers CD44 is the most documented CSC marker presented within HNSCC patient samples^27-29^. CD44 was found to be a viable target of the Wnt pathway. For the maintenance of stemness of CSCs, Wnt pathway is considered as a main pathway and also accepted as a poor prognostic marker^30^.

Clinical findings stipulate that CSCs exhibit intrinsic property of resistance to therapy. They can vigorously adjust to different environmental conditions, e.g., hypoxia and lack of nutrients. Though the fundamental molecular mechanisms are still not well understood, but it was corroborated that CSCs manifest an increased scavenging of reactive oxygen species (ROS), enhanced capacity of DNA repair, reduced induction of apoptosis and enhanced cell survival which play major role in therapy resistance and promotes poor patient"s outcome^31,32^. This resistance causes treatment failure in various types of cancer. Therefore, to identify and to target CSCs is indispensable to improve the success rate of treatment^33^.

Sun et al (2010)^34^ and Lu et al (2011)^35^ have observed that patients exhibiting high expression of CD44 have greater tendency to develop lymph node metastasis, recurrence and resistance to radiotherapy. In non-Hodgkin"s lymphoma and colon cancer, over expression of CD44 has been correlated with advanced stage and poor survival^36^. In addition to this, Wang et al (2006)^37^ and Kawano et al (2004)^38^ observed association of CD44 with lymph node metastasis, tumor volume and poor survival in HNSCC. In renal cell carcinoma, an association between CD44 and stage, grade and poor prognosis was observed^39^. Collectively, these studies stipulated that presence of CD44 in a tumor tissues or cells might play a key role in treatment failure, lymph node metastasis, poor survival and recurrence.

Along with CD44 and other TNM factors, HPV/p16 positivity is also a known risk factor and an independent prognostic factor. However, this has been excluded from our study because our study is mainly focused on determining the independent predictive significance of CD44. If HPV positivity was taken into account it may confound our results particularly in specimens exhibiting positivity for both CD44 and HPV infection. The average age for the diagnosis of HNSCC ranging between 55-65 years^40^ whereas in our study majority of patients were of age < 45 years. Sharma et al (2015)^41^ have reported that in their study 46.2% patients were belonged to 31-50 years of age group. Also, in Indian population the consumption of tobacco (smokeless and tobacco smoke) and alcohol is quite prevalent, particularly in Northern India people start consuming alcohol and tobacco at a very early age. This explains our finding that in our study majority of patients were of <45 years.

In various previous studies, the role of CD44 and its isoforms especially CD44v has been established in the prognosis and progression of HNSCC. But, current study was mainly focused on the assessment of CD44 expression and no isoform of CD44 has been studied because the involvement of CD44 in response to radiotherapy and OS in HNSCC had not been delineated. Our findings are consistent to the previous studies showing profound clinical importance of CD44. An attempt can be made to understand the clinical relevance of CD44. Our data suggest that patients with high expression of CD44 can increase the number of CSCs that are mainly resistant to radiotherapy which leads to recurrence and tend to have shorter survival. Although, the overall response rate to therapy in our study is low as compared to previous studies. Owing to the fact that in our study, the majority of the patients were of stage IV with N2 lymph node metastasis and T4 tumor size exhibiting more aggressive tumors particularly in younger patients. Additionally, high positivity of CD44 which is also attributed to chemo-radioresistance is also accountable for low overall response rate. Our results corroborate the findings in previous studies establishing that presence of CD44 was correlated with treatment failure, lymph node metastasis, advanced stage, tumor size, grade, recurrence and poor survival. We can also infer through our findings that except CD44 there could be other parameters that could serve as viable targets for the better treatment such as alcohol consumption, lymph node metastasis and grade. An evident aspect of current study was that CD44 may be an indicator of poor prognosis in advanced stage patients of HNSCC. Therefore, based on our findings, we can propose that it may be necessary to employ all the viable targets in order to determine the effective treatment modalities and to gather all the relevant information which can be transcribed into patient benefit.

In conclusion, our study demonstrated that high CD44 expression, positive lymph node status, high grade and advanced stage are the predictive markers for treatment and OS which contributes to treatment failure, poor survival and recurrence. However, further exploration is warranted to understand the mechanism of CD44 including all other isoforms of CD44. The ratio of CD44 along with other markers of CSCs could also be evaluated in forthcoming studies to understand the mechanism of CSCs in treatment failure which will further help clinicians to devise an effective anticancer therapy of improved efficacy.

## Acknowledgement

We are thankful to the Vice Chancellor of KGMU and the Director of CDRI for providing the facility and infrastructure. We are also thankful to all the patients for their participation in the study.

## Author’s Contribution Statement

Conception and design: MadanLal Brahma Bhatt and Parul Dubey

Analysis and interpretation of data: MadanLal Brahma Bhatt and Parul Dubey

Drafting, review and/or revision of the manuscript: MadanLal Brahma Bhatt, Parul Dubey, Smrati Bhadauria, Rajeev Gupta, Anupam Mishra, and Vijay Kumar

Final approval: MadanLal Brahma Bhatt and Smrati Bhadauria

