## Supplementary data for "Evaluation of Correlation between CD44, Radiotherapy Response and Survival Rate in Patients with Advanced Stage of Head and Neck Squamous Cell Carcinoma (HNSCC)"

**Suppl 1: Different primary tumor sites of HNSCC (n=90)**

| **Tumor Sites** | **No. of Patients**  **n=90 (%)** |
| --- | --- |
| **Oral Cavity:**  Alveolus  Base of Tongue  Border of Tongue  Buccal Mucosa  Hard Palate  Oral Cavity  Retromolar Trigone  Soft Palate  Tongue  Tonsil  Tonsillar Fossa | 4 (4.4)  10 (11.1)  18 (20.0)  20 (22.2)  1 (1.1)  3 (3.3)  1 (1.1)  1 (1.1)  7 (7.8)  8 (8.9)  1 (1.1) |
| **Pharynx:**  Hypopharynx  Oropharynx  Pyriform Fossa | 4 (4.4)  2 (2.3)  1 (1.1) |
| **Larynx:**  Glottis  Larynx  Supraglottic Larynx | 1 (1.1)  2 (2.3)  3 (3.3) |
| **Nasal Cavity:**  Nasal Cavity  Nasopharynx | 1 (1.1)  2 (2.3) |

**Suppl 2: Demonstrates percentage of frequency distribution of CD44 expression in primary tumors of different sites in HNSCC (n=90)**

| **Location** |  | **CD44 Expression** | | **Total** |
| --- | --- | --- | --- | --- |
|  |  | **Low** | **High** |  |
| Oral Cavity | Count  Percentage | 16  21.6% | 58  78.4% | 74  100% |
| Pharynx | Count  Percentage | 1  14.3% | 6  85.7% | 7  100% |
| Larynx | Count  Percentage | 0  0.0% | 6  100% | 6  100% |
| Nasal Cavity | Count  Percentage | 0  0.0% | 3  100% | 3  100% |
| Total | Count  Percentage | 17  18.9% | 73  81.1% | 90  100% |

The majority of tumor specimens were collected from oral cavity (n=74) whereas n=7, 6 and 3 tumor specimens were collected from pharynx, larynx and nasal cavity respectively. Furthermore, majority of patients were exhibited high expression of CD44.
